# Synthetic Substitutes as a Conservation Tool: Evaluating Synthetic Leopard Fur for Demand Reduction and Species Recovery

**DOI:** 10.1101/2025.07.31.666914

**Authors:** Aditya Shekhar Malgaonkar, Gareth Whittington-Jones, Tristan Dickerson, Maswabi Lishandu, Sarah Davies, Zoe Woodgate, Xia Stevens, Choolwe Mulenga, Gift Mulenga, Mathew Phiri, Lucky Mulenga, Hon. Manyando Mukela, Gryton Kasamu, Willem A. Nieman, Gareth Mann, Abishek Harihar, Diogo Veríssimo, Rob Pickles

**Affiliations:** Panthera, New York, New York, USA; Barotse Royal Establishment, Mongu, Western Zambia; Wildlife Crime Prevention, Lusaka, Zambia; Institute for Communities and Wildlife in Africa, University of Cape Town, Cape Town, South Africa; Panthera Wild Cat Conservation Zambia Limited, Lusaka, Zambia; Department of National Parks and Wildlife, Chilanga, Lusaka, Zambia; Biodiversity and Behavioral Science Team, Environmental Change Institute, University of Oxford, Oxford, UK

## Abstract

Providing synthetic substitutes is a widely promoted strategy to shift consumer demand away from wildlife products derived from threatened species. Yet, there is little evidence on whether such product substitution interventions effectively prevent illegal or unsustainable harvesting and contribute to the recovery of threatened populations. Drawing on the Furs For Life (FFL) Zambia initiative, which supplied synthetic furs known as “Heritage Furs” to replace leopard skins traditionally worn during Lozi royal ceremonies in Western Zambia, we present an evaluation designed to test both the effects and causal mechanisms of substitution. Guided by the EMMIE framework, commonly used in crime prevention evaluation, we triangulated data from semi-structured questionnaires, law enforcement patrols, court records, camera trap monitoring, and stakeholder interviews conducted between 2018 and 2024. Qualitative analysis using the General Elimination Method was employed to assess plausible alternative explanations for leopard recovery. By 2024, adoption of synthetic furs among leopard fur users exceeded 80 percent, while self-reported ownership of authentic leopard furs declined by 70 percent. At the same time, patrol detections of leopard poaching incidents decreased, and camera trap density estimates increased from an average of 2.7 to 3.8 leopards per 100 square kilometers across the focal landscape. An integrated mechanism of change, derived from stakeholder perspectives, indicates that while substitution reduced demand, concurrent and reinforcing effects of counter-poaching and counter-trafficking operations were critical to leopard recovery. This study provides the first empirical link between a demand reduction initiative based on synthetic substitutes and measurable species population recovery.

## INTRODUCTION

Illegal and unsustainable wildlife trade is widely recognized as a leading proximate threat to global biodiversity, accelerating population declines and extinctions across taxa (Morton et al., 2021; Hinsley et al., 2023). In a meta-analysis of 506 sites spanning 145 vertebrate species, populations exposed to threats were, on average, 60% smaller than matched controls, and ∼16% were locally extirpated (Morton et al., 2021). Annual revenues to transnational illegal wildlife trade networks are estimated at US$ 7–23 billion per year (UNODC 2020), with ecological costs borne by species valued for wild meat, traditional medicine, ornamentation and the live-animal trade.

Conventional, enforcement-centered responses such as international trade bans, protected-area designations and law enforcement patrols have slowed losses in some areas but often fail to halt declines (Geldmann et al., 2013; Challender & MacMillan, 2014; Hsiang & Sekar, 2016). Moreover, many biodiversity hotspots lie in lower-income nations where wildlife agencies are chronically under-staffed and under-resourced (Dutton et al., 2013). Under these conditions, prohibitions can inflate black-market prices, attract organized crime and displace hunting into remoter areas (Armsworth et al., 2020; Hsiang & Sekar, 2016). Furthermore, coercive approaches may also alienate Indigenous Peoples and Local Communities (IP&LCs) who depend on wildlife for nutrition, income or cultural identity, and thus undermine legitimacy and effectiveness of conservation efforts (Challender & MacMillan, 2014; Torrents-Ticó et al., 2023).

To address these shortcomings, product substitution strategies based on the provision of high-fidelity synthetics or alternative natural products that satisfy user needs without wild harvest have gained significance (Broad & Burgess, 2016). If substitutes are affordable, accessible and culturally acceptable, they can reduce the scarcity premium that fuels poaching (Armsworth et al., 2020; Thomas-Walters et al., 2021). Additionally, if a substitute conveys key symbolic cues such as prestige or ritual authenticity, users face minimal switching costs and behavioral change becomes more likely (Rock & MacMillan, 2022). Recent examples and emerging case studies are encouraging. In India, 3-D-printed tiger teeth and hornbill casques are being trialed for ceremonial regalia (Dhar, 2023). A survey of 2463 Vietnamese consumers showed a marked preference for synthetic bear bile over farmed or wild bile (Davis et al., 2022). A discrete-choice experiment with 300 shark-fin consumers in Singapore found stronger willingness to buy certified-responsible or lab-cultured fins than conventional wild-caught fins (Choy et al., 2024). In South Africa, the Furs for Life project halved leopard-skin use among Nazareth Baptist Church congregants over ten years (Naudé et al., 2020). Despite these signs of promise, robust evaluations remain scarce, leaving managers uncertain about the magnitude, durability and possible unintended effects of substitution strategies.

The evidence gap is emblematic of a broader shortfall in counter-wildlife-crime research. A systematic review of field-based measures to curb poaching and trafficking assessed 123 studies, but identified only five with evaluation designs capable of attributing causation, none of which tested a product-substitution intervention (Delpech et al., 2021). Gaps are most acute where interventions intersect with IP&LC ceremonial practices, which often impose stringent authenticity requirements (Torrents-Ticó et al., 2023). To date, peer-reviewed studies of synthetic substitutes have focused on consumer attitudes, stated intentions or adoption rates (cf. Naudé et al., 2020; Davis et al., 2022; Choy et al., 2024); rigorous evaluation of conservation outcomes such as reduced harvesting or population recovery is almost entirely absent (Rytwinski et al., 2021).

Evaluators therefore must ask whether interventions (i) curb consumption, (ii) measurably diminish direct threats to targeted species populations and (iii) promote stable or increasing population trends, while also (iv) illuminating the causal pathways that link actions to outcomes. This is challenging because illegal trade is covert, baseline data are sparse, and multiple interventions often overlap (Baylis et al. 2016). Quantitative indicators such as snare detections, seizure volumes and density estimates show what changed, but seldom why (Johnson et al., 2015). Moreover, seizures, for instance, may rise simply because enforcement effort increased (Paudel et al., 2022).

Evaluation designs integrating qualitative insights from interviews, focus groups and document analysis with quantitative data are increasingly advocated to connect observed changes to underlying mechanisms (Mascia et al., 2014; Zavaleta-Cheek et al., 2023). The EMMIE framework (Effect, Mechanism, Moderator, Implementation, Economic cost) provides a structured template for such evaluations (Johnson et al., 2015). For example, a recent assessment of counter-poaching patrols in Malaysia linked specific tactics (Mechanism) to reductions in snaring (Effect) and identified landscape and offender characteristics that moderated success (Lam et al., 2023). While EMMIE broadens the scope of enquiry, it does not specify how to infer causality. The General Elimination Method (GEM) complements EMMIE by systematically ruling out rival explanations until the most plausible causal narrative remains (Scriven, 2008; Salazar, 2019). Embedding GEM within the Effect–Mechanism–Moderator steps yield a transparent chain of reasoning, particularly valuable in wildlife-trade contexts where randomized or perfectly matched controls are rarely feasible and where extrinsic effects can confound attribution (Baylis et al., 2016).

The ceremonial use of leopard (*Panthera pardus*) furs and resultant leopard poaching in western Zambia illustrates both the challenge and the opportunity. Each year the Barotse Royal Establishment (BRE) and Lozi People mark the flood cycle of the Barotse Floodplains with the Kuomboka (outbound) and Kufuluhela (return) ceremonies. During the main Kuomboka, around 200 royal paddlers ferry the Litunga - the King of the Lozis on a black-and-white ‘Nalikwanda’ barge while wearing red berets trimmed with lion mane and ‘Lipatelo’ skirts comprised of leopard, serval (*Leptailurus serval*), genet (*Genetta* spp.), civet (*Civettictis civetta*) and, rarely, cheetah (*Acinonyx jubatus)* furs. Leopard furs - symbolizing power, courage and grace are preferred, while other spotted carnivore furs are alternatively used in the absence of leopard fur or added to enhance appearance. An estimated 1000 prospective paddlers vie for paddler places each year, generating demand for an estimated 150–250 wildlife fur garments annually (Panthera, 2022a).

Paddlers obtain leopard skins via two channels: (i) self-hunting, and (ii) poacher-to-paddler trade in which middle-men commission local hunters or paddlers buy directly from hunters. All such activity is illegal without a permit under the Zambian Wildlife Act (2015). In the Greater Kafue Ecosystem (GKE) - a landscape comprising of Kafue National Park and surrounding Game Management Areas, the Department of National Parks and Wildlife (DNPW) partners with Panthera, African Parks, Game Rangers International and Musekese Conservation to conduct counter-poaching patrols and counter-trafficking interventions. While offenders caught hunting or transporting furs may be arrested, law-enforcement agencies refrain from seizing garments at the Lozi ceremonies out of cultural sensitivity. Interviews with key community contacts, questionnaires with paddlers and seizure records confirm that the GKE is the principal source of ceremonial leopard skins (Panthera, 2022a). Poaching and trafficking of leopard furs is believed to be tightly linked with the ceremonial fur demand (Whittington-Jones et al., 2020).

To stem this pressure, the late Senior Chief HRH Inyambo Yeta invited Panthera in 2017 to partner with the BRE to replicate South Africa’s Furs for Life model for the leopard poaching problem in Western Zambia. The resulting project, initially named Saving Spots, now Furs for Life (FFL) Zambia, began distributing high-fidelity synthetic leopard skins in 2019. Implementation (2019– 2024) coincided with intensified patrols and trafficking operations and overlapped with COVID-19 lockdowns, creating a complex setting in which multiple interventions, deep-rooted cultural norms and external shocks interact.

Our study is the first rigorous evaluation of a wildlife-product substitution strategy that tracks outcomes along the full supply chain—from consumer demand, trafficking and poaching to population trends of the target species. By analyzing multiple data sources and explicitly accounting for contextual complexity, the study provides evidence that synthetic substitutes can curb illegal trade and drive measurable species recovery.

## METHODS

### Evaluation approach

We used the EMMIE framework to structure our evaluation, integrate multiple evidence streams and link intervention delivery to ecological outcomes. To strengthen causal inference, we nested two complementary tests within this scaffold: Eck’s Four-Point Test (Eck, 2017) to determine causality and the General Elimination Method (GEM) to investigate alternative explanations and develop an integrated mechanism for change (Figure 1).

**Figure 1.**
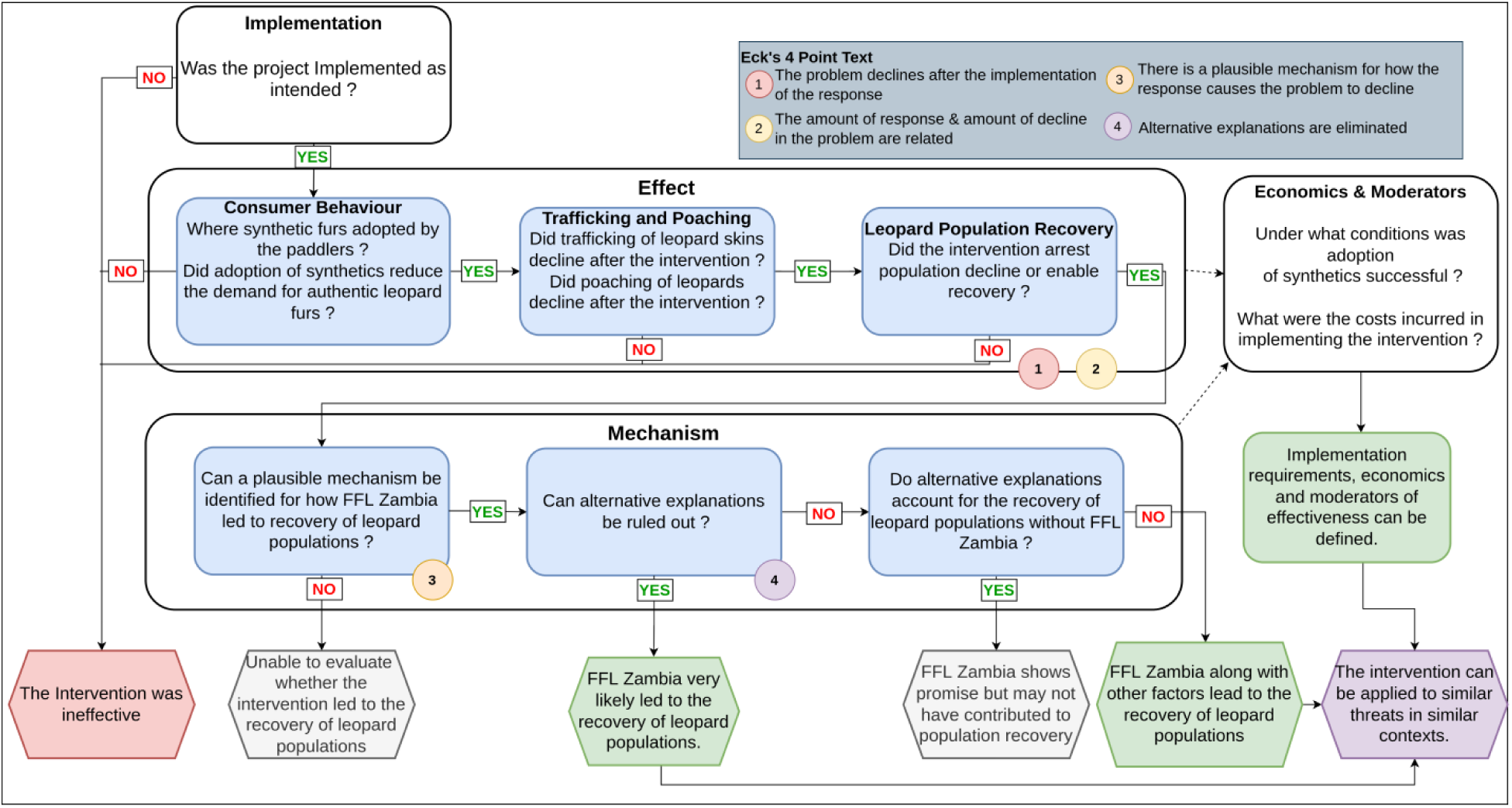
Workflow documenting the EMMIE evaluation approach used in this study.

First, we verified that the synthetic-fur roll-out was implemented as planned, documenting any challenges and adaptations; this established the factual basis for analyzing effect and mechanism. Next, we quantified intervention effects at three nested levels; (i) consumer demand for leopard furs, (ii) trafficking/poaching pressure, and (iii) leopard population trend, thereby tracing the complete causal chain predicted by the project logic. At this stage, Eck’s test allowed us to judge whether change followed implementation and whether the magnitude of change matched the scale of the intervention. We then applied GEM to determine whether a credible mechanism linked the intervention to the observed changes and if alternative explanations could account for observed outcomes – thereby fulfilling criteria for Eck’s four-point test for causality.

Finally, we identified *Moderators*, i.e. context-specific factors that facilitated synthetic-fur adoption and downstream leopard recovery and addressed the *Economics* dimension by estimating total project costs and expressing cost-effectiveness as the cost per leopard-use prevented.

#### 1. Implementation

We assessed if FFL Zambia was implemented as designed by tracking project implementation starting from the issuing of royal declaration and synthetic fur distribution to the use of synthetic furs at Kuomboka ceremonies. The implementation roll-out was reconstructed from project reports, and the impact of delays caused due to disruptions such as the Covid-19 pandemic and adaptation measures used were documented.

#### 2. Effect

##### 2.1 Consumer behavior change, and change in leopard fur use

A preliminary survey was conducted in 2018 to establish a baseline, during which two researchers recorded the number of leopard, serval, and other carnivore furs worn by paddlers at the Kuomboka ceremony. Following project initiation, four structured questionnaire surveys targeting Lozi royal paddlers or prospective paddlers (2020: *n* = 86; 2021: *n* = 23; 2022: *n* = 57; 2024: *n* = 110) documented their (i) awareness of synthetic furs, (ii) perceptions of synthetic furs, (iii) intention to acquire authentic leopard furs, (iv) current fur ownership, and (v) use of authentic leopard furs at the Kuomboka ceremony. The questionnaires were administered in the Lozi language by a local Lozi researcher (ML) familiar with cultural context and practices (see *Supplementary Material S1*).

For each year, we used exact binomial tests to examine whether observed proportions for each of the five binary response variables (i–v) differed significantly from a random expectation of 0.50 and reported associated 95% confidence intervals and *p*-values (Sokal & Rohlf, 2012). To assess temporal changes in the same binary response variables, we fitted generalized linear models (GLMs) with a binomial error distribution and a logit link function, treating “year” as a continuous predictor. Regression coefficients (β) and their associated *p*-values were used to evaluate temporal trends in awareness, perception, intention, ownership, and use (Crawley, 2013). Model assumptions were assessed using residual diagnostics and visual checks for overdispersion. Statistical significance for all tests was accepted at *a* = 0.05. All analysis was conducted using R statistical software version 4.4.1 (R Core Team, 2024).

##### 2.2 Trafficking and poaching trends

We used information from Wildlife Crime Prevention (WCP) Zambia’s court-monitoring program covering wildlife crime related cases across 16 courts in western and southern Zambia (1069 case records; 2018–2024), and law enforcement patrol data collected from the GKE, (196,600 km foot patrols; 2018–2022) to index trafficking and poaching, respectively. Poaching detections from GKE were examined descriptively due to the limited data available.

##### 2.3 Leopard density trends

Twenty-four camera trapping surveys were conducted in five sectors across the GKE, between 2018 and 2024. To investigate temporal changes in leopard density (individuals/100 km^2^), we applied maximum likelihood multi-session spatial capture-recapture (SCR) methods (Borchers & Efford, 2008), treating each year as a separate session. SCR models estimate population density (*D*), by incorporating the spatial locations of individual detections to estimate baseline detection probability (*g0*) and the spatial decay parameter (*σ*) (Gerber & Parmenter, 2015). While SCR is typically used to estimate density at a single point in time, multi-session (i.e., multi-year) data can be modelled to describe population trends (Royle et al., 2013). In multi-session SCR models, parameters, such as g0 and σ, are shared across sessions/years, smoothing excess sampling variance and increasing model accuracy (Rogan et al., 2019). To account for sex-specific capture heterogeneity (Sollmann et al., 2011), all models included sex as a covariate on σ and were modelled using a partially observed finite mixture. All SCR models were fitted using the package *secr* 5.2.4 (Efford, 2025). As each session represented a ‘closed’ population, we specified the total area of leopard habitat sampled per session, by creating a buffer around each station equal to four times σ. We restricted this model state space to avoid modelling density within urban areas or water bodies (Plummer et al., 2017).

We estimated the finite growth rate, or population trend, as the change in leopard density between sessions (λ*_t_*), by reparametrizing the density model such that:

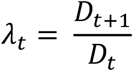

where *D_t_* represents density at year *t* (Efford, 2025; Hadi et al., 2025).

#### 3. Mechanism and alternative explanations

To identify the mechanisms through which demand-reduction might have contributed to leopard recovery in the GKE, we carried out 22 semi-structured interviews with purposively selected informants from three stakeholder groups central to the poaching issue: Lozi community leaders and paddlers, DNPW officers, and conservation-NGO staff (see *Supplementary Material S2*). Interviews were conducted in person or via Microsoft Teams™ between February and April 2024, in English or, for community members, in Lozi with an interpreter. After informed verbal consent, each session began with background questions, followed by a ’leopard population trend-exercise’. Respondents were shown a graph of leopard-density estimates for 2018–2022 and a deck of twelve cards, each naming a potential causal factor derived from global leopard assessments (Jacobson et al., 2016; Stein et al., 2023), regional case studies (Panthera, 2022a; Panthera, 2022b) and field experience of authors (GWJ, TD). Interviewees sorted the cards into: (A) factors believed to have positively influenced leopard numbers, (B) factors not contributing positively, and (C) unsure/unknown, and could add new factors (see *Supplementary Material S2*). They then ranked Category A factors from most to least influential while explaining their reasoning; probing questions explored causal pathways and interactions. Card order was shuffled between interviews. The exercise was replicated with digital cards in a shared Miro® workspace for remote sessions. Rankings and narratives were recorded in a field notebook; factor rankings and respondent metadata were collated using a spreadsheet, and narratives were transcribed into a text document.

Narrative data were manually coded to standardize recurring phrases and iteratively identify themes. We combined each interviewee’s response into frequency scores for each potential causal factor, stratified scores by stakeholder group, and assessed the supporting rationale provided by interviewees. Following the General Elimination Method (Scriven, 2008; Salazar, 2019), we built a stakeholder group-specific mechanism of change that retained any factor cited by at least one-third of respondents in that group. The three group-specific mechanisms were merged: factors endorsed by more than one group were retained, and pathways were refined for logical coherence. The resultant integrated mechanism of change was cross-validated against information from patrol records captured using the Spatial Monitoring and Reporting Tool (SMART), wildlife-crime court cases, and case studies (Panthera, 2022a; Panthera, 2022b). Expected outputs implied by the integrated mechanism were compared with observed trends to assess the plausibility of proposed mechanisms linking synthetic fur adoption, complementary interventions, and extrinsic events to reduced poaching pressure and leopard recovery.

#### 4. Moderators effectiveness

Questionnaire responses from paddlers were stratified based on their opinion on synthetic furs, desire to acquire authentic furs, and use of authentic furs at Kuomboka, and respondent covariates (age, experience, employment, and education status) were inspected to identify contextual moderators of synthetic fur adoption.

#### 5. Economics of implementation

We estimated the cost per adult leopard use prevented by dividing the total project expenditure by the estimated number of whole leopard skins no longer required for ceremonial garments as a result of the intervention. Project expenditure comprised three components: audited start-up costs incurred during the initiation phase (2018–2020); actual maintenance outlays over the first five fiscal years (2020–2024); and the net present value of projected maintenance costs for the subsequent five years (2025–2029), discounted at 8% annually to account for high local inflation.

To estimate the number of whole leopard skins prevented from ceremonial use, we applied a Fermi-estimation approach that decomposed the calculation into a sequence of observable or estimable parameters. These included: the annual number of prospective paddlers participating in the Kuomboka ceremony; the pre-intervention proportion wearing authentic leopard skins; the lifespan of authentic leopard-skin garments; the number of garments derived from a single leopard; the ten-year lifespan of synthetic furs; and the annual adoption rate of synthetic garments. These inputs were used to estimate annual demand for leopard skins and the number prevented each year due to substitution. Summing the annual number of whole leopard skins prevented over the 2020– 2029 evaluation window yielded the total number of adult leopards prevented from ceremonial skin use (For parameter values see *Supplementary Material S3*).

Where possible, we applied mean or median, minimum, and maximum values for each parameter to generate three scenarios: likely, conservative, and optimistic and used them to calculate corresponding scenario-based cost-per-leopard estimates. To enable cost comparison with area-based conservation measures – for which costs are documented in published scientific literature, we estimated the area equivalent by multiplying the number of leopards prevented annually by ‘100/baseline leopard density per 100 km^2^’, this represents the area these animals would otherwise occupy. Cost per unit area (USD/km²) was calculated by dividing the total intervention cost by the area equivalent.

## RESULTS

### 1. Implementation assessment

Consultations with the Barotse Royal Establishment-the traditional Lozi royal authority-facilitated the design of highly realistic and culturally appropriate synthetic fur garments, known as “Heritage Furs” modelled on authentic leopard and serval fur patterns. The initial batch of Heritage Furs were unveiled at the official public launch of FFL Zambia in August 2019 and used for the first time by 70 royal paddlers accompanying the Litunga during a *Kupuwana* Ceremony held in September 2019. Following feedback, subsequent batches of Heritage Furs were redesigned, and by early 2022, 1,350 synthetic fur garments were delivered to the BRE – which was responsible for distribution of furs prior to the ceremonies, retrieval, and storage.

In December 2019, the Litunga issued a royal decree mandating that only synthetic furs would be worn in future official Lozi ceremonies, framing the adoption of synthetics as a way to uphold tradition while protecting wildlife. This was communicated via a radio broadcast by the BRE’s Prime Minister and reiterated during a ceremony in November 2019.

The COVID-19 pandemic resulted in the cancellation of ceremonies in 2020 and 2021, delayed fabric shipments, and limited in-person community engagement, posing a risk to sustained interest. The project adapted by shifting outreach online, using WhatsApp messaging and YouTube videos to promote synthetic furs with positive, tradition-centered messaging.

Synthetic furs were distributed for use at the 2022 Kuomboka ceremony and again at the 2024 ceremony following the cancellation of the Kuomboka in 2023. A local outreach officer maintained engagement with the community, conducted surveys, and distributed educational materials, while a project coordinator managed logistics and communication with the BRE and Panthera. Senior traditional leaders played a vital role in sustaining cultural acceptance and legitimacy.

### 2. Effect

#### 2.1 Consumer behaviour change, and change in leopard fur use

Across the four survey (2020–2024), awareness of synthetic furs was consistently high with 81.9 % of prospective paddlers (226 of 279) being aware of them, a proportion significantly greater than random expectation (p < 0.01; 95 % CI = 78–100 %). Although awareness dipped in 2021, it increased significantly between 2021 and 2024 (β_year_ = 0.51, p < 0.01; Figure 2). Attitudes mirrored this pattern with 78.5 % of respondents (215 of 276) expressing a favorable opinion of synthetic furs (exact binomial, p < 0.01; 95 % CI = 73–100 %). Following a transient decline in 2021, approval climbed to 94.4 % in 2024 (β_year_ = 0.95, p < 0.01).

**Figure 2.**
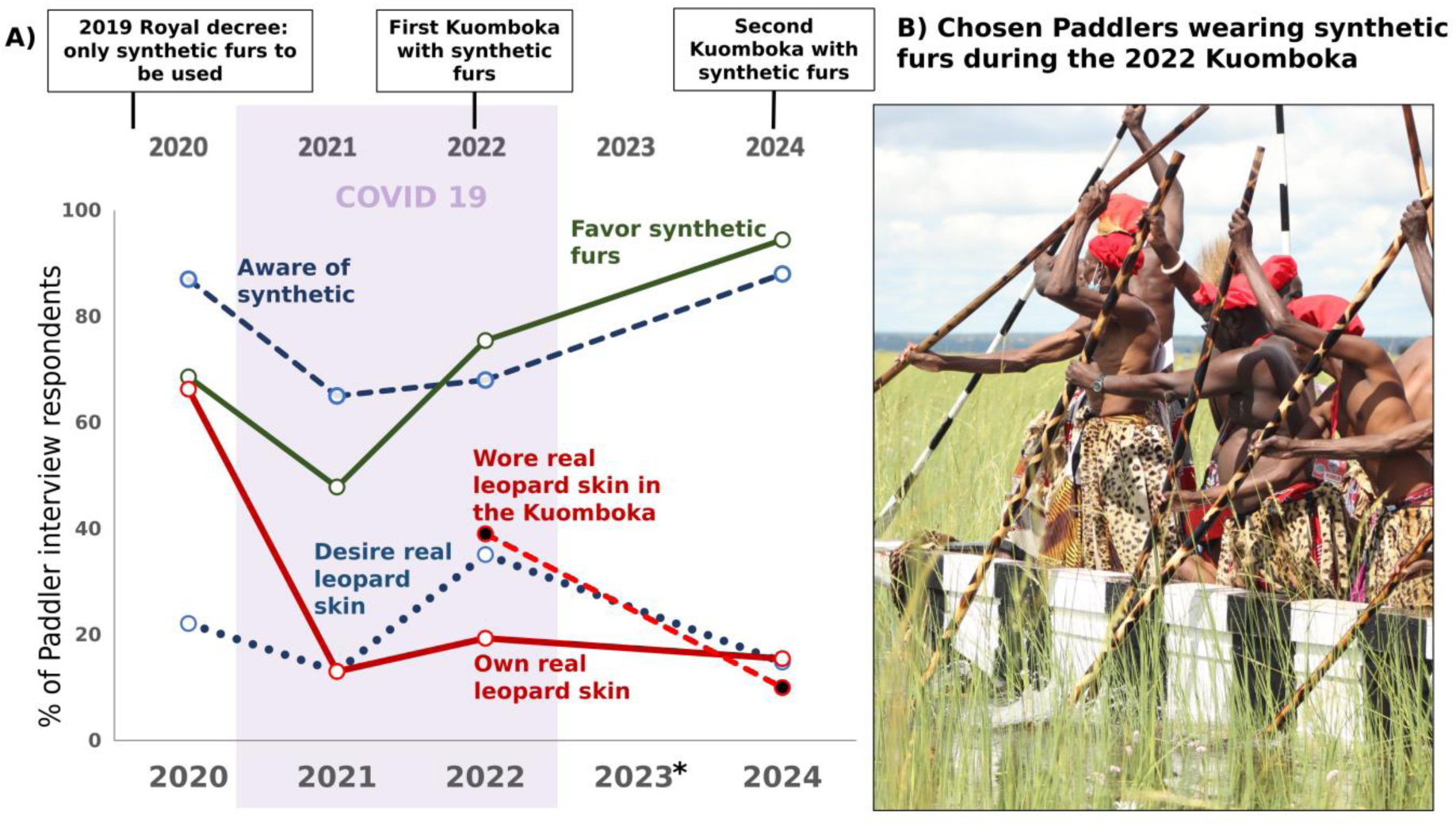
Change in perceptions among prospective paddlers from interview surveys in relation to key events in the FFL Zambia intervention (A), and an example of synthetic furs worn by royal paddlers during the 2022 Kuomboka. * No paddler interview surveys were conducted in 2023.

The desire to acquire authentic leopard furs remained low and stable. Only 21 % of paddlers wished to obtain authentic fur, significantly below the 50 % benchmark (p < 0.01; 95 % CI = 17– 26 %), and this proportion showed no temporal trend (β_year_ = –0.10, p = 0.25). Actual ownership, however, declined sharply (Figure 2). In 2020, 67% of paddlers (57 of 86) possessed at least one leopard fur; by 2024, this had declined to 15%, representing a 77.6% reduction in ownership over the four years (β_year_ = –0.59, p < 0.01). Correspondingly, use of authentic furs during the Kuomboka ceremony plummeted (β_year_ = –1.69, p < 0.01) with 39% of royal paddlers wearing authentic leopard furs in 2022, compared with just 10% in 2024, representing a 74.4% decline over two years (Figure 2).

#### 2.2 Trafficking and poaching trends

The court monitoring recorded 1,069 wildlife-related arrest and seizure incidents from 2018 to 2024. The number of leopard-related seizures declined significantly from 11.25 per 100 wildlife crime cases in 2018 to 2.88 per 100 cases in 2024 (β_year_ = –0.30, p = 0.02). However, the data does not suggest that seizures of other spotted carnivores declined during the same period (β_year_ = 0.12, p = 0.432).

Seven leopard poaching incidents were reported from GKE in 2019, four of which were recorded by patrols, and three incidents were recorded through court monitoring data (Figure 3). In 2020, the number of reported poaching incidents did not change, with six incidents recorded through court monitoring and one through ranger patrols. However, just one incident of leopard poaching was reported in 2021, and, in the following years up to 2024, no incidents of leopard poaching have been reported.

**Figure 3.**
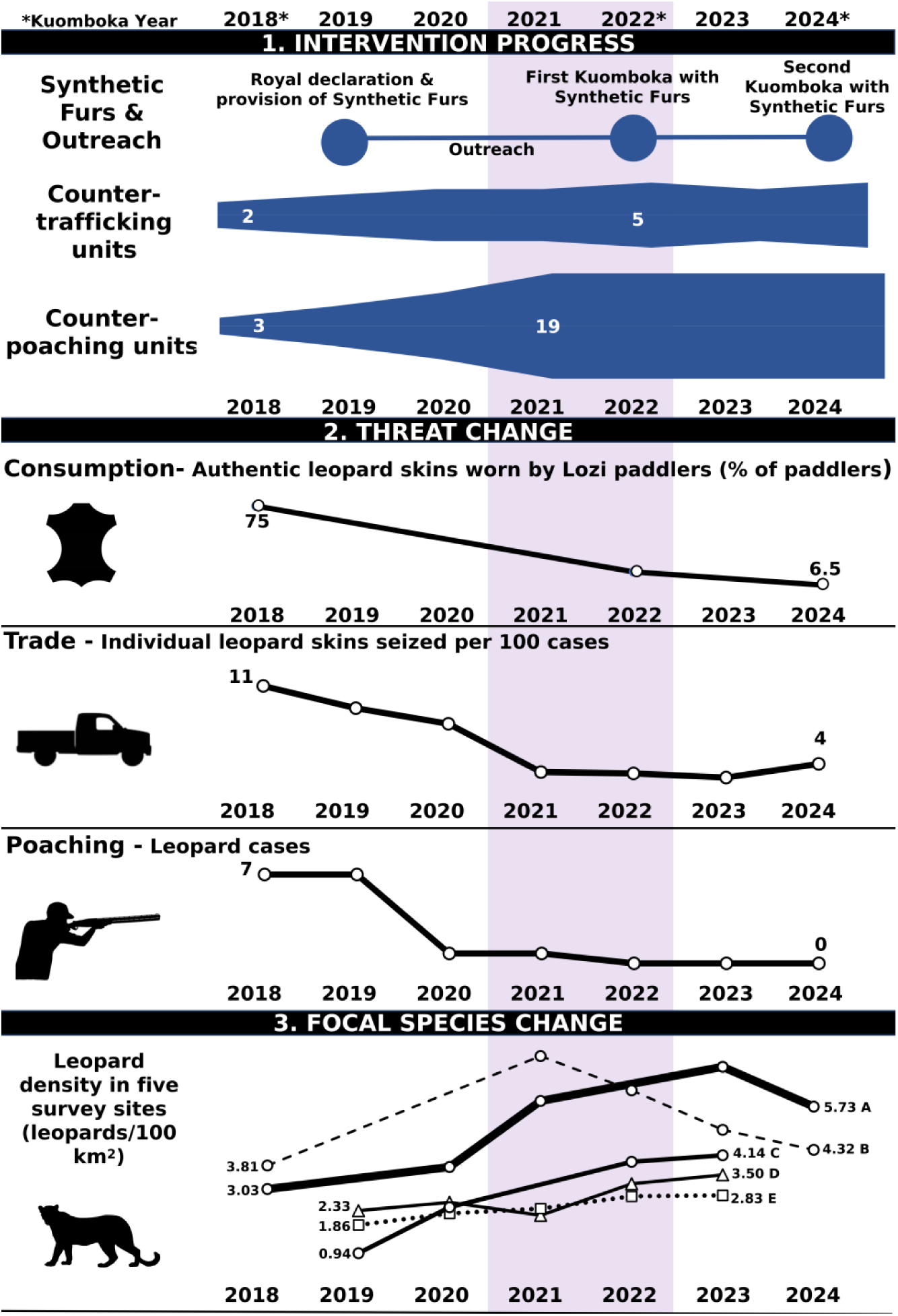
Evolution of interventions, declines in leopard trade, and rise of focal leopard population in Kafue National Park in four survey sites (A= Kafue Central; B= Mumbwa GMA; C= Mulobezi GMA; D= Kafue North, E = Kafue South)

#### 2.3 Leopard density trends

Estimates from multi-session SCR models (Figure 3), as well as λ_overall_ (i.e., mean λ*_t_* for each sector; Table 2), indicated leopard population growth across the GKE (with λ_overall_ >1.0 for most sectors). Estimated leopard densities were higher than ’baseline estimates’ (i.e., estimated density derived from the first year of camera trapping). These sites exhibited linear population growth, with little to no fluctuations in annual density estimates. While λ_overall_ was greater than 1 for both GMAs (Table 2), multi-session SCR models highlighted an apparent downturn in population growth in leopard density for Mumbwa (after a peak in 2021; Figure 3). Notably, Mumbwa’s 2024 density estimate did not significantly differ from the ’baseline density’ (4.32/100 km^2^ [95% CI: 3.24 – 5.76] versus 3.81/100 km^2^ [95% CI: 2.73 – 5.31], respectively).

**Table 1:** Eck’s Four Point causality test for FFL Zambia project.

| Test | Evidence and Conclusion |
| --- | --- |
| Decline in the problem comes after the intervention | Pass. Following the implementation of the FFL Zambia project, a rapid and sustained decline in demand for authentic leopard furs was observed. Interview data revealed a 78% reduction in authentic fur ownership among paddlers between 2020 and 2024, with reported desire for authentic furs declining concurrently.<br>Law enforcement data further corroborated these trends. Recorded poaching incidents across the Greater Kafue Ecosystem declined from a total of 14 incidents recorded during 2018 to 2019, before the intervention, to zero incidents recorded from 2022 to 2024. Court-monitoring records similarly show a 75% decrease in leopard fur seizures during the same period (2018–2024), suggesting a marked disruption of the trafficking supply chain.<br>In parallel, Leopard population densities increased in all five monitored sectors from 2018 to 2024. Notably, two sectors demonstrated a twofold increase in density, rising from approximately 2 to 4 individuals per 100 km<sup>2</sup>. The temporal alignment between the onset of the FFL Zambia intervention and these demographic improvements, coupled with concurrent reductions in poaching and trade indicators, provides strong inferential support that the intervention contributed to demand reduction and population recovery outcomes for leopards in the GKE. |
| The amount of intervention and the amount of the problem's decline are related. | Pass. Adopting synthetic furs leads to a proportionate decline in leopard fur ownership. Decline in fur ownership, similarly, resulted in a proportionate decline in the trafficking of leopard furs. However, the reduction in poaching-related mortality of leopards could not be quantified. |
| A clear mechanism by which the intervention caused the decline | Pass. A clear mechanism linked the implementation of the FFL Zambia project, with reduction in authentic fur use and ownership, trafficking of furs, and poaching of leopards in GKE. The adoption of synthetic furs, reinforced by a royal decree and community outreach, significantly reduced ceremonial demand for authentic leopard furs, collapsing the local market for poached furs, reducing poaching incentives, and enabling leopard population recovery across the Greater Kafue Ecosystem in conjunction with parallel supply-side interventions. |
| Alternative explanations are rejected. | Partially Pass. While intensified counter-poaching patrols and anti-trafficking operations likely contributed to reduced leopard mortality, the scale and timing of leopard recovery suggest that these efforts alone were insufficient to account for the observed outcomes. Notably, leopard population recovery began in two of the five sectors before the adoption of SMART-enabled patrols but shortly after the launch of the FFL Zambia intervention in 2019, which included the royal directive banning authentic furs and the rollout of synthetic alternatives. Moreover, the consistent and significant decline in leopard fur ownership and ceremonial use, over 70% between 2020 and 2024, coincided with widespread adoption of synthetic furs and was not preceded by similar trends in enforcement activity alone. These patterns align more strongly with mechanisms of demand reduction than with traditional supply-side enforcement. |

**Table 2:**
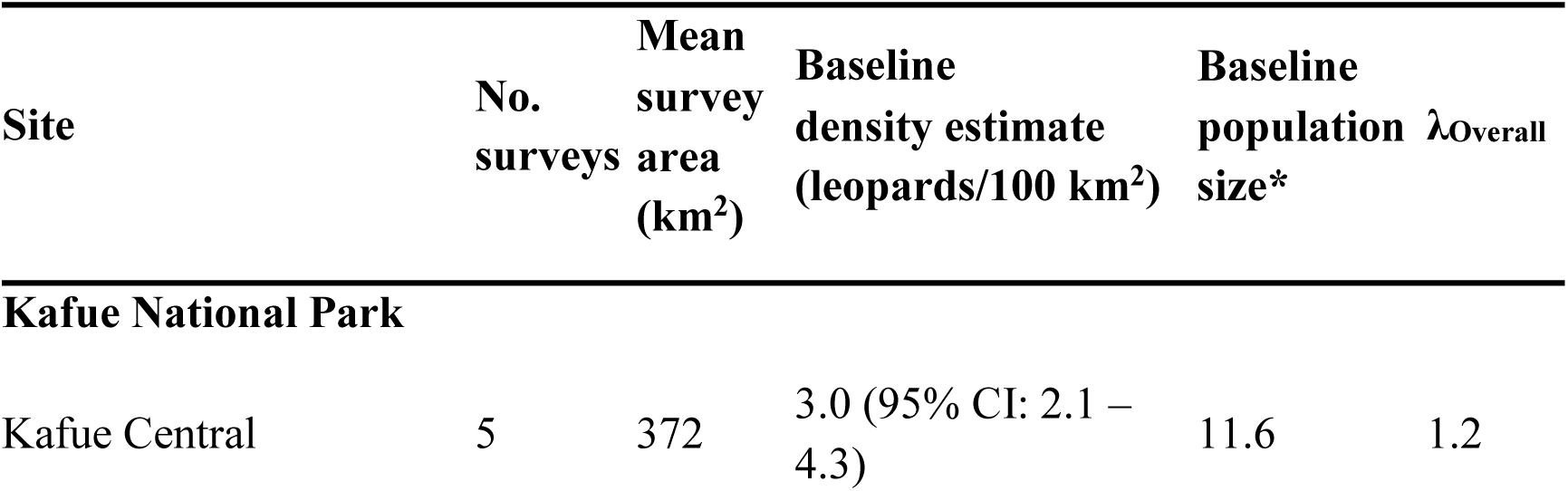
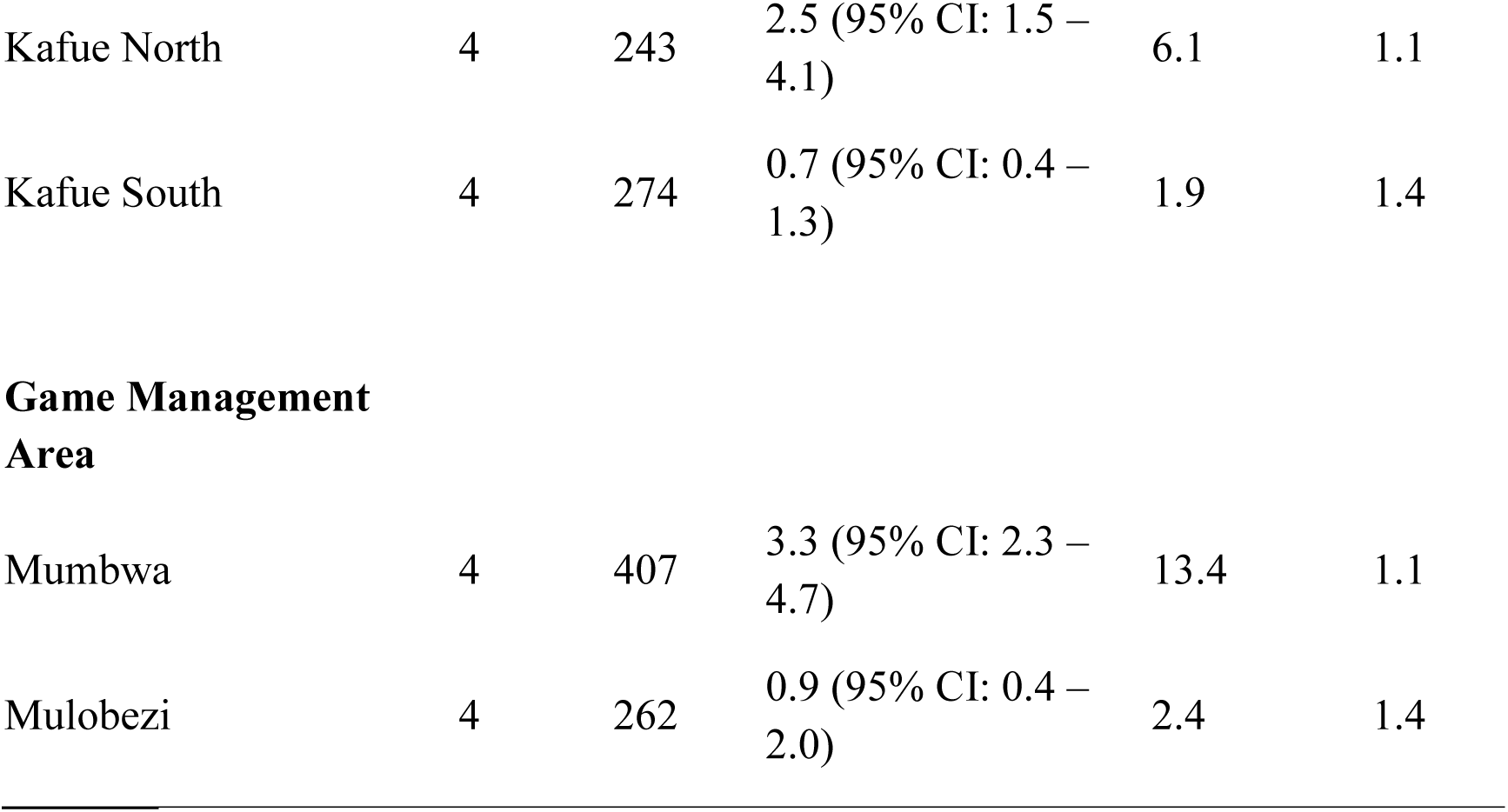
Overall population growth (λ_Overall_) for sites with leopard density estimates derived from multi-session spatial capture recapture (SCR) models.*Baseline population size was estimated by multiplying the estimated density (per km^2^) from the first camera-trap survey undertaken at each surveillance site by the mean survey area of that site.

### 3. Mechanism and alternative explanations

#### 3.1 Demand for leopard fur reduced due to synthetic furs

Sixteen of the 22 stakeholders we interviewed, including all 10 Lozi community representatives and 6 of 7 conservation-NGO staff, ranked FFL Zambia as the first or second most influential driver of the leopard recovery in the GKE. Interviewees agreed that the switch from authentic furs to synthetic furs reduced demand for leopard furs.

Community stakeholders stressed four attributes of synthetic fur that, in their opinion, accelerated adoption: they were (i) readily available, (ii) distributed free of charge, (iii) visually appealing, and (iv) socially acceptable to the wider community (A1, A3–A5). Additionally, community stakeholders linked the difficulty in procuring authentic leopard furs to the rapid adoption of substitutes (A1, A3–A5). Interviewees stated that providing synthetic furs and conservation messaging fostered a cultural shift favoring leopard conservation (B4, B7, A9). This shift likely limited resistance from traditionalist sections of society and facilitated the wide adoption of synthetic furs (A4, A5, A6, A9, B2).

All of the community and most conservation NGO interviewees (6 of 7) opined that reduced market demand likely weakened the motivation for targeted poaching. A few interviews (A6, A8, A10) suggested that demand reduction impacted paddler-driven targeted poaching and poaching by local poachers for sale to traders and paddlers. No interviewee suggested that synthetic fur availability had stimulated “fashion” demand outside the Lozi community, countering a common substitution risk.

#### 3.2 Counter-trafficking operations caused market disruption

Nineteen of the 22 interviewees representing all three stakeholder groups identified counter-trafficking operations as a key causal factor for leopard recovery. Support was unanimous among conservation-NGO stakeholders (7 of 7), and high among community (8 of 10) and DNPW stakeholders (4 of 5). Interviewees reported that the operations heightened the legal risks faced by offenders, disrupted trafficking networks, and removed prolific offenders through arrest and conviction (A5, A6, B3, B4, B6, C1–C3). By dismantling these networks, the interventions were thought to restrict poachers’ opportunities to market leopard furs, lowering the financial incentive for killing leopards (B2, C2, C3).

Heightened probabilities of detection, arrest, and prosecution were also cited as major deterrents to poaching (B2, B6, C1–C3). Respondent B6 noted that visible court follow-ups and publicized convictions further increased perceived risk among potential offenders. The incapacitation of key offenders through imprisonment was viewed as reducing ongoing poaching pressure on leopards and their prey (B4, B6, C3). Overall, interviewees perceived that elevated legal risks, disruption of trade networks, and the removal of key traffickers were critical to curbing illegal activity and supporting leopard population recovery.

#### 3.3 Counter-poaching patrols reduced opportunity and increased perceived risks

Counter-poaching patrols were considered one of the key drivers of leopard recovery by 16 of 22 interviewees across all three stakeholder groups. Patrols were ranked the top intervention by most DNPW respondents (3 of 5) and conservation-NGO staff (5 of 7). Interviewees emphasized that wider patrol coverage and higher patrol frequency improved detection of illegal activities, including leopard and prey poaching (B2, A8, B7, B5, C1–C4). Hiring local community scouts was viewed as pivotal, strengthening community support, information flow, and the timely reporting of offences (B7, B4).

Greater detection raised the perceived likelihood of arrest and prosecution; together with the incapacitation of likely prolific offenders, this reportedly deterred would-be poachers (A3, A9, B2, B3, B4, C1–C5). Patrol teams also directly disrupted poaching by seizing snares and firearms and regularly interrupting illegal activities (C1–C4). These actions increased the costs and effort required to poach, reducing its feasibility. Overall, heightened detection, increased legal risk, and higher operational costs for offenders were seen as central to the decline in leopard and prey poaching, facilitating leopard population recovery across the GKE.

#### 3.4 Integrated Mechanism of Change

A clear causal mechanism is evident with culturally endorsed substitutes eliminating ceremonial demand, disrupting the illegal market, and reducing poaching incentives. While counter-trafficking and counter-poaching contributed to the recovery of leopard populations by disrupting trafficking and bushmeat poaching in GKE (Figure 4). For a summary of available evidence supporting the causal pathways see *Supplemantary Material S4*.

**Figure 4:**
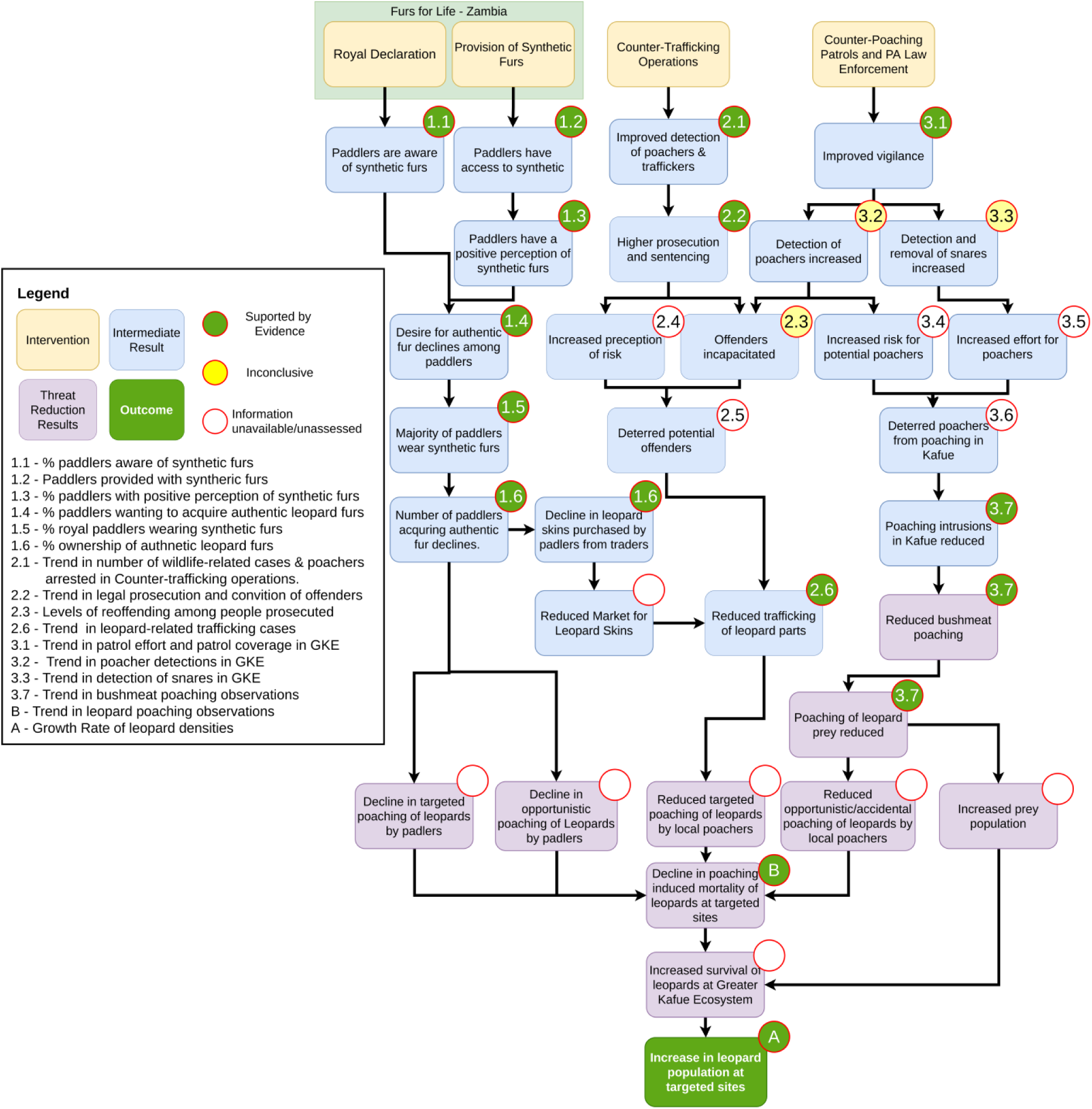
Integrated mechanism of change illustrating validated pathways for interventions leading to leopard recovery and evidence rating.

The Eck’s four-point test indicates strong evidence of the effectiveness of the FFL Zambia project (Table 1). The decline in leopard fur ownership, trafficking, and poaching occurred after the intervention began in late 2019, with the scale of the decline proportionate to the extent of synthetic fur adoption. Thus, the intervention meets three criteria fully and one partially, providing robust inferential support for its impact on leopard recovery.

### 4. Moderators of effectiveness

The interviews identified two distinct categories of fur users among paddlers: facultative and adherent fur users. The overall proportion of prospective paddlers expressing a negative opinion of synthetic furs was 14% (39 of 276); however, 67% of them wanted to acquire authentic leopard furs in the future compared to just 16% of the paddlers with a favorable opinion of synthetic furs (p < 0.01). In 2020, for which information is available, 53% of prospective paddlers with negative opinions admitted to hunting leopards for their furs. There was no evidence to suggest that level of education (p = 0.32) or employment status of respondents (p = 0.30) influenced the desire to acquire new leopard furs. However, 85% of prospective paddlers with a negative opinion (n = 56) reported the need for preserving traditions and culture as a reason for their opinion of synthetic furs. Conversely, among those with a favorable opinion (n = 215), quality and realism of synthetic fur were cited as a reason by 52%, while 26% cited conservation of leopards and other wildlife. This suggests further decline in the acquisition of leopard furs may be limited by ’adherent fur users’ who primarily cite the need to preserve traditions as a motivation.

### 5. Economics of Implementation

The total cost of implementing the leopard-skin substitution intervention over a ten-year period (2020–2029) was estimated at USD 691,678. This comprised an initial start-up expenditure of USD 313,558, incurred between 2018 and 2020 for infrastructure, community consultation, and design, production, distribution and promotion of synthetic fur. Recurring maintenance costs including staff salaries, community engagement, repairs, and monitoring are projected at USD 378,120 over the 10-year implementation window.

Our analysis estimated that, in the absence of intervention, approximately 176 authentic leopard-skin garments would be required each year to meet ceremonial demand among prospective paddlers. Based on the average yield of four garments per leopard, this translates to roughly 44 leopards killed annually. Over the ten-year period following the introduction of synthetic alternatives, and accounting for the rate of adoption and the longevity of synthetic garments, the intervention is estimated to have prevented the use of 360 whole adult leopards (36 leopards/year), with a plausible range of 204 to 1310 leopards under conservative and optimistic scenarios, respectively. When compared against the total program cost, the cost per leopard use prevented was calculated at approximately USD 1,924 in the likely scenario of preventing the use of 360 leopards, USD 528 for the optimistic scenario (1310 leopards) and USD 3,395 under the conservative estimate of 204 leopards prevented. Considering the mean baseline leopard density of 2.7/100 km^2^, this is equivalent to USD 69167.8 spent each year to conserve a leopard population spread over an area of 1333.33 km^2^, translating into USD 51.88/km^2^ per year.

## DISCUSSION

Our evaluation suggests that the FFL-Zambia project reduced demand for authentic leopard skins and contributed to species recovery by providing culturally acceptable synthetic furs, with backing from traditional leadership and support from complementary enforcement efforts.

### Synthetic substitutes can reduce demand and support species recovery

Quantitative results from our paddler surveys revealed a >70% decline in authentic leopard-fur ownership between 2020 and 2024, far exceeding the ∼40–50% reduction expected solely from effective supply restriction. This sharp decline coincided with the provision of culturally endorsed synthetic furs, suggesting a targeted effect rather than a general market constraint. Reported trafficking of leopard furs fell by over 50%, and average leopard density estimates from the GKE increased by approximately 65%, indicating population-level recovery. Increases in leopard density in Kafue Central, Kafue North, Mumbwa GMA, and Mulobezi GMA followed the rollout of SMART-enabled patrols and the launch of the FFL Zambia initiative after the royal declaration in late 2019. However, leopard recovery in Kafue South and Mulobezi GMA preceded SMART-enabled patrol deployment, indicating that observed trends were not solely due to increased enforcement.

Stakeholder interviews supported the role played by synthetic furs in demand reduction. Acceptance of synthetic furs was driven by social legitimacy, visual similarity to authentic furs, free access, and endorsement by the Barotse Royal Establishment. These findings are consistent with broader research showing that product substitutes are more likely to succeed when they fulfill functional, aesthetic, and symbolic needs (Thomas-Walters et al., 2021; Broad & Burgess, 2016; Rock & MacMillan, 2022). Similar dynamics have been observed in China, where declines in shark-fin consumption followed strong social endorsement and near-indistinguishable alternatives (Jeffreys, 2016; Zhou et al., 2021). Studies on lab-grown rhino horn also highlight that perceived authenticity and trusted leadership endorsements are critical for uptake (Chen, 2017).

### Product-substitution using synthetics is an economically viable approach

To our knowledge, this is the first study to report a cost-effectiveness estimate for demand-reduction interventions. Preventing the use of one adult leopard cost approximately USD 1,924, with comparable annualized area-equivalent cost of USD 52/km². This is significantly lower than the USD 7,133 per animal reported for rhino dehorning in South Africa (Kuiper et al., 2025) and falls well below standard benchmarks for effective protected area management in Africa, typically ranging from USD 138 to USD 2,000/km²/year (Lindsey et al., 2018; Green et al., 2012; Blom et al., 2004). While this comparison is not exact as protected areas support many species and ecosystem services, and our estimate reflects only ceremonial demand for leopards, it offers a valuable perspective and suggests that product substitution using synthetic alternatives can be an economically viable strategy.

### Effectiveness of substitutes amplified by complementary interventions

Although substitution likely played a central role, its effects were amplified by complementary interventions. Intensified counter-poaching patrols raised the perceived risk of detection and limited poaching opportunities. Counter-trafficking operations and prosecution efforts disrupted supply chains by increasing the risk of arrest and potentially removing key offenders. Together, these enforcement actions likely created a deterrent effect that was reinforced by collapsing consumer demand. These findings reflect broader literature emphasizing that supply-side enforcement and demand reduction can be more effective when implemented together (Challender & MacMillan, 2014; Veríssimo & Wan, 2019). Demand reduction alone cannot address residual poaching driven by alternative markets or prestige-seeking users, while enforcement alone often displaces rather than eliminates illegal activity unless end-user demand also declines.

Stakeholders consistently cited outreach and awareness initiatives including community meetings, radio programs, and social media messaging as key to raising risk perception and community buy-in. This supports findings from behavioral research, which emphasize that interventions combining credible messaging, norm reinforcement, and visible deterrents achieve stronger compliance (Thomas-Walters et al., 2021; Thomas-Walters et al., 2023).

Our results underscore that successful substitution strategies require integration with supply-side controls, community engagement, and leadership backing to address inertia, cultural attachment, and organized trade.

### Structured frameworks integrating qualitative insights can enhance evaluations

A key strength of this evaluation was the use of the EMMIE framework and the General Elimination Method to integrate quantitative and qualitative information. EMMIE enabled us to link observed effects to underlying mechanisms, implementation context, moderating factors, and economic costs. The General Elimination Method helped differentiate between plausible alternative explanations, enhancing confidence in causal inference. This is especially important in demand-reduction contexts, where changes in consumer behavior may also result from supply constraints. For example, increased adoption of synthetic fur and reduced trafficking of leopard skins could be attributed to heightened park-level enforcement. However, interviews revealed that most paddlers adopted synthetic furs voluntarily and for conservation reasons, suggesting a demand-side mechanism.

While qualitative methods have long been recommended in conservation evaluation (Salafsky, 2002), few studies apply them rigorously in interventions involving IP&LCs. In such situations, where collective norms, deeply rooted beliefs and traditional authority shape behavior, qualitative insights are crucial to understand how an intervention is received and endorsed by communities (Torrents-Ticó et al., 2023). By capturing paddlers’ perspectives on tradition, social acceptance, and emerging norms, we gained deeper insight into both successes and resistance among more conservative segments.

We were unable to estimate leopard poaching rates or determine cause-specific mortality directly. Additionally, while we focused on ceremonial demand among paddlers, we did not examine broader patterns of leopard use or potential shifts in bushmeat consumption that could influence prey availability. Monitoring prey density alongside predator recovery would help clarify these relationships, especially in prey-limited systems (Creel et al., 2018). These gaps reflect the inherent challenges of evaluating conservation interventions in complex settings where direct attribution is rarely straightforward. Nonetheless, by drawing on diverse stakeholder perspectives, assessing alternative explanations, identifying key assumptions, and tracing plausible causal pathways, this evaluation aligns with the core principles of robust qualitative attribution (Zavaleta-Cheek et al., 2023).

## CONCLUSION

Our study demonstrates that culturally endorsed synthetic substitutes, when combined with community engagement and partnerships, traditional leadership, and supply-side controls, can reduce demand for illegal wildlife products and support species recovery. With costs comparable to conventional supply-side measures, synthetic substitution offers an economically viable option. The use of EMMIE and the General Elimination Method to link observed effects to plausible explanations and trace causal pathways shows the value of combining qualitative and quantitative evidence in conservation evaluation. Together, our findings position culturally grounded product substitution using synthetic replicas as an effective, economically viable and complementary addition to the conservation toolkit.

## ACKNOWLEDGEMENTS

We gratefully acknowledge the Lozi People, Barotse Royal Establishment, the Department of National Parks and Wildlife, the National Prosecution Authority, and Wildlife Crime Prevention (WCP) Zambia and Panthera colleagues past and present for their collaboration and support. We are especially indebted to the late Senior Chief of the Lozi people, His Royal Highness Inyambo Yeta (1954 to 2023), whose vision and leadership were pivotal to the inception and success of the FFL Zambia initiative. We also thank Musekese Conservation, the Zambian Carnivore Programme, African Parks, Game Rangers International, and Kashikoto Conservancy for their valuable contributions to the project and ongoing conservation efforts in the Greater Kafue Ecosystem. We thank Dr. Kim-Overton Young for the guidance and support provided for this study and Dr. Andrew Lemieux for his guidance for the initial Saving Spots evaluation. Finally, we are grateful to our donors: Cartier for Nature, Peace Parks Foundation, the Royal Commission of AlUla (Saudi Arabia), and the UK Department for Environment, Food and Rural Affairs for their generous support.

## ETHICS STATEMENT

The interviews and questionnaire surveys conducted for this study was approved by the Research Ethics Committee, Faculty of Science, University of Cape Town, in May 2019. Reference No: FSREC 92 - 2019.

## USE OF ARTIFICIAL INTELIGENCE TOOLS

We used OpenAI ChatGPT (model-04) to check for grammar, spelling errors and refine language to improve readability. The authors have reviewed and edited the content as needed and take full responsibility for the content of this manuscript.

