## Supplementary Material 1 for "Synthetic Substitutes as a Conservation Tool: Evaluating Synthetic Leopard Fur for Demand Reduction and Species Recovery"

**Supplementary Material S1: Paddler Questionnaire**

|  | **Question** | **Response Options** |  |  |  |  |
| --- | --- | --- | --- | --- | --- | --- |
|  |  | A | B | C | D | E |
| **Basic Background Details** | Are you willing too participate in this survey? | Yes | No |  |  |  |
|  | Interview Date | [Date Feild) |  |  |  |  |
|  | What is your home location (grid reference on map) | [Grid on Map] |  |  |  |  |
|  | What is your age? | [Numeric] |  |  |  |  |
|  | What is your highest level of education | None | Primary | Secondary | Tertiary |  |
|  | What has been your predominant emplacement status during the previous 5 years? | Unemployed | Employed | Partially Employed |  | Word of Month |
|  | What media do you most commonly consume | Newspaper | TV | Social Media | Radio |  |
| **Participation at Kuomboka, wild cat skin preference and use** | Have you ever been a paddler in the Kuomboka? |  |  |  |  |  |
|  | For how many years? | (Numeric Field) |  |  |  |  |
|  | What is the significance of spotted cat skins for the Lozi? | (Descriptive) |  |  |  |  |
|  | Which species is more prestigious? | Leopard | Cheetah | Serval | Genet | Other (text) |
|  | Which cat skin (photo) do you prefer? | Cheetah | Genet | Leopard | Serval |  |
|  | When do the Lozi start loking for skins in preparatio for the Kuomboka? | (Text) |  |  |  |  |
|  | How often do you replace your skins? Every XXX years | (Numeric Field) |  |  |  |  |
|  | How do you store your skins and/or prevent damage to them? | (Text) |  |  |  |  |
|  | Which cat skins do you own ? | Leopard | Cheetah | Serval | Other |  |
|  | How many leopard skins do you own? | (Numeric Field) |  |  |  |  |
|  | How many cheetah, serval, or other species skins do you own? |  |  |  |  |  |
| **Leopard skin acquisition** | How did you obtain these leopard skins? | Inherited | Purchased | Hunted | Other(text) |  |
|  | Where did you purchase the leopard skin? | (Text) |  |  |  |  |
|  | Did you purchase a full leopard skin, half a skin, or strips? | Full | Half | Strips | Other(text) |  |
|  | How much did the leopard skin cost? | (Numeric Field) |  |  |  |  |
|  | Did the person have a leopard skin on hand or did you have to place an order? | (Text) |  |  |  |  |
|  | What hunting method do you use for leopard? | Shooting with Firearm | Wire snare | Foot trap | Other(text) |  |
|  | Where do you typically go hunting for leopard? | (Map Grid) |  |  |  |  |
|  | How many cheetah, serval, or other species skins do you own? |  |  |  |  |  |
| **Opinion of Heritage Furs** | Where did you first hear about the Heritage Furs? |  |  |  |  |  |
|  | What is your opinion of the Heritage Furs? ( and Why) |  |  |  |  |  |
|  | Do you intend to get a new wild cat skin in the future? Why? |  |  |  |  |  |
