## Supplementary Material 2 for "Synthetic Substitutes as a Conservation Tool: Evaluating Synthetic Leopard Fur for Demand Reduction and Species Recovery"

Supplementary Material S2:

I. Interviewees representing the three stakeholder groups

| **Stakeholder Group** | **Code** | **Organisation** | **Occupation/Position** |
| --- | --- | --- | --- |
| Community | A1 | - | Businessman/ Paddler |
|  | A2 | - | Fisherman/Paddler |
|  | A3 | - | Community Leader/ Retd. Military Official |
|  | A4 | - | Community Leader/Businessman |
|  | A5 | - | Community Leader/Paddler |
|  | A6 | - | Community Leader/Tourist Guide |
|  | A7 | - | Businessman/Paddler |
|  | A8 | - | Tailor/Paddler |
|  | A9 | - | Journalist |
|  | A10 | - | Tailor/Paddler |
| Conservation NGOs | B1 | Panthera | Community Engagement Officer |
|  | B2 | Panthera | Leadership Position |
|  | B3 | WCP | Leadership Position |
|  | B4 | WCP | Leadership Position |
|  | B5 | Musekese Conservation | Community Engagement Officer |
|  | B6 | Zambian Carnivore Programme | Researcher/ PhD Scholar |
|  | B7 | Panthera | Leadership Position |
| Department of National Parks and Wildlife (DNPW) | C1 | DNPW | Ranger/Wildlife Police Officer |
|  | C2 | DNPW | Ranger/Wildlife Police Officer |
|  | C3 | DNPW | Leadership Position/Senior Police Officer |
|  | C4 | DNPW | Ranger/Wildlife Police Officer |
|  | C5 | DNPW | Ranger/Wildlife Police Officer |

II. Predefined Potential Causal Factors
1. Anti-poaching patrols in the Greater Kafue Ecosystem
2. Anti-trafficking operations targeting trade in illegal wildlife parts/products
3. Education and Awareness Campaigns conducted by conservation NGOs
4. Saving Spots Program: Synthetic Heritage Furs provided for Lozi Ceremonial Gatherings
5. Covid-19 pandemic and its after-effects
6. Increase in wildlife and eco-tourism in Protected Areas and Game Management Areas within the last decade.
7. Increase in employment in Zambia within the last decade.
8. Decrease in local demand for leopard parts/products for Traditional African Medicine.
9. Decrease in local demand for wild meat/bushmeat.
10. Enhanced regulation of trophy hunting in Game Management Areas.
11. Improved habitat management practices in Protected Areas and Game Management Areas.
12. Reduction in Human-Leopard Conflict in and around Protected Areas and Game Management Areas.
III. Stakeholder Interview Guide

Interviewer Statement and Respondent Consent

Hello, I am ____________ I work with a Wildlife Conservation NGO and we are conducting a survey to understand how conservation interventions has affected leopard populations in Greater Kafue Ecosystem. As part of this survey, I would like to interview you and document your opinions through a questionnaire and simple card exercise. Your answers and responses to the exercise will be documented and used for research. The information and results generated through this study will be used for internal and publicly available reports, and for a scientific article. You may choose to keep your identity anonymous.

Will you like to participate in this survey?

Will you like us to keep your identity anonymous and confidential?

Do you have any queries with regard to this survey ?

Will you like us to contact you with further questions/additional surveys in the future?

Section 1: Basic Background Information

Full Name:

Gender and Age Group:

Contact Details:

Official Designation/Occupation :

Agency:

Date and Time of Interview :

Location of Interview :

1. Can you describe your current role and field of expertise/field of work ?
2. How long have you been in this role ?
3. Do you or your organization contribute to wildlife conservation ? If so, How ?
4. What areas of western Zambia have your worked in ? (geographic area)
5. Have you heard of Panthera’s Saving Spots Program or Synthetic Heritage Leopard Furs ?

Section 2 : Leopard Population Trend Exercise (Present the Graph Slide to the Respondent)

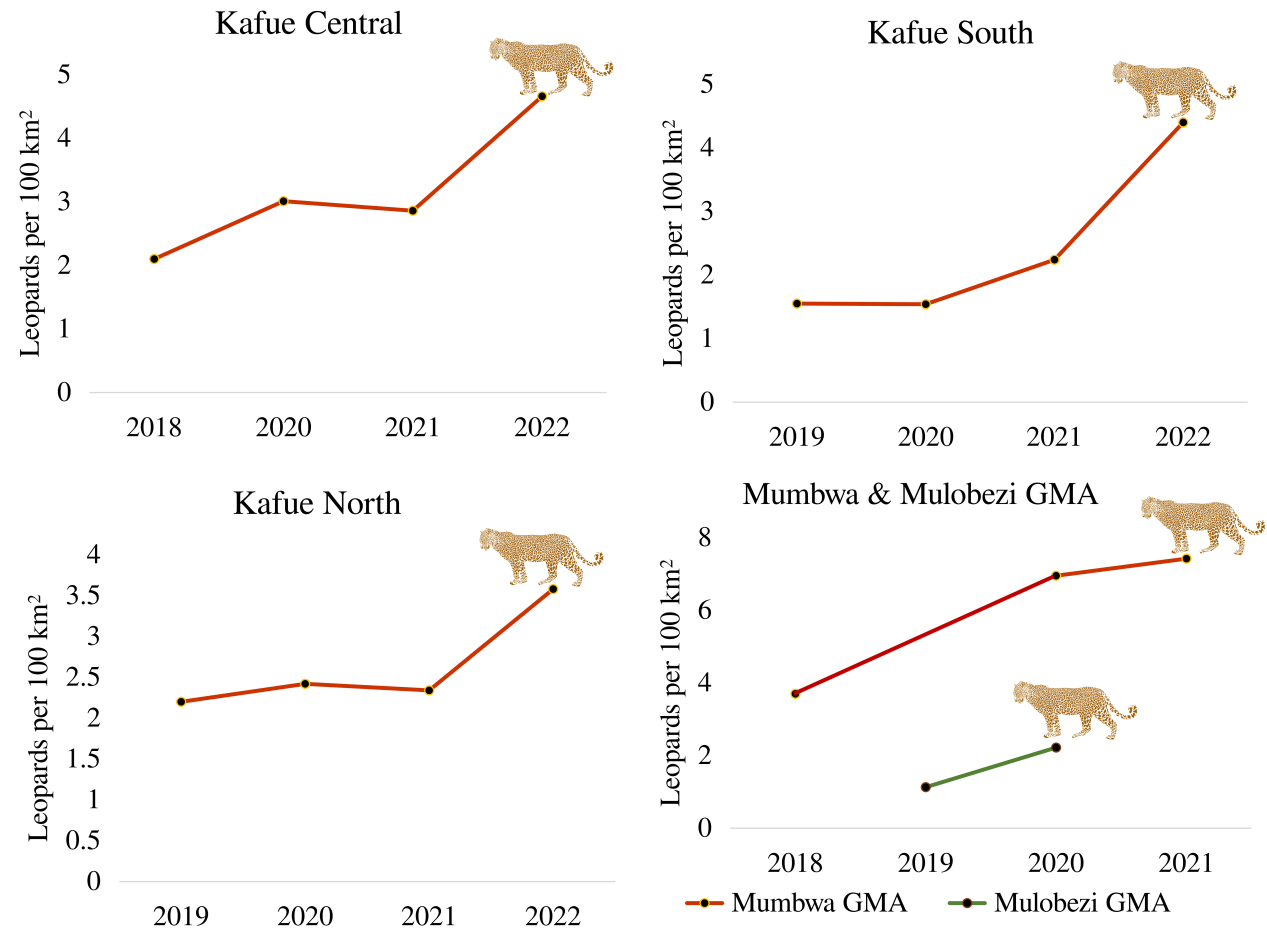

*“The graphs shows the trend in leopard populations as measured by estimates of adult leopard per 100 sq. kms in Greater Kafue Ecosystem from 2018 to 2022. Please take a moment to examine the graphs.”*

1. Do you agree that leopard densities in Greater Kafue Ecosystem have increased since 2018 as depicted in the above graphs ?
2. If No, Do you think leopard populations are stable or declining, why you think so ?
3. Can you place the following cards in three piles ? A) Factors that have affected the leopard population since 2018 B) Factors that have not affected the leopard population since 2018 C) Unsure/Unknown.
4. In addition to these factors, do you think there are any other factors which could be responsible for the rise in leopard densities ? Place the factor in Pile A (Write down new factors, place them in the pile)
5. Can you now sort the cards in Pile A form the most influential factors affecting leopard densities to the least influential factors ? ( Left to right from most influential to the least) (Capture Multiple Photos)
6. Can you explain why do you think each of these factors have affected leopard densities ?
7. Can you explain how factor X led to the observed trend in leopard densities ?
