## Supplementary Material 3 for "Synthetic Substitutes as a Conservation Tool: Evaluating Synthetic Leopard Fur for Demand Reduction and Species Recovery"

**Supplementary Material S3: Fermi Estimation Parameters for Cost-per Leopard use prevented.**

| A | B | C | D | E | F | G | H | I | J |
| --- | --- | --- | --- | --- | --- | --- | --- | --- | --- |
| Sr. No | Value | Mean Estimate | Minimum | Maximum | Calculations | Information Source/Remarks |  |  |  |
| A1 | Estimated proportion of royal paddlers(%) wearing authentic leopard skin at Kuomboka before 2020 (pre introduction of synthetics) | 0.75 | 0.75 | 0.9 | Input assumption | Preliminary field survey in 2018. Questionnaire survey of prospective paddlers and field observation in 2020 (pre‑synthetic) |  |  |  |
| A2 | Minimum number of paddlers seeking to participate in Kuomboka each year | 1000 | 1000 | 1250 | Input assumption | Expert opinion and key community contacts. |  |  |  |
| A3 | Mean estimated longitivity of authentic leopard skin garments ( in years) | 4.25 | 2 | 5 | Input assumption | 2020 paddler questionaire; median durability |  |  |  |
| A4 | Estimated longitivity of synthetic leopard furs (in years) | 10 |  |  | Input assumption | Field inspection of 2019 synthetic garments |  |  |  |
| A5 | Estimated mean no. of garments made from a single dead leopard | 4 | 2 | 6 | Input assumption | Field observations & Photo‑analysis of whole skins vs paddler skirts |  |  |  |
| B1 | How many new authentic leopard skin garments are required each year ? | 176.470588235294 | 150 | 562.5 | (A2 × A1) ÷ A3 | Annual demand for new authentic leopard skin garments |  |  |  |
| B2 | How many dead leopards/whole skins are required to supply authentic fur garments for Kuomboka each year ? | 44.1176470588235 | 25 | 281.25 | B1 ÷ A5 | Leopards potentially killed per year to supply demand |  |  |  |
| B3 | Initial Project Start-Up Cost(2018-2020) in USD | 313558 |  |  |  | Project financial records (2018–2020), real 2025 USD |  |  |  |
| B4 | Project Maintainance Cost ( 2020 onwards) in USD | 37589.26 |  |  |  | Based on 2022–23 audited accounts; deflated by 8% per year for 2 years, escalated at C5 thereafter |  |  |  |
| C1 | Discount rate (real, per year) | 0.08 |  |  | Input assumption | discount rate accounting for high inflation rate in Zambia (~ 14 %) |  |  |  |
| C2 | Annual replacement/failure rate of synthetic garments | 0 |  |  | Input assumption | Attrition can be mitigated through the surplus skins supplied. |  |  |  |
| C3 | Synthetic Fur adoption rate Year 1 (proportion using synthetics) | 0.65 |  |  | Input assumption | Adoption in first season post‑rollout 2020 |  |  |  |
| C4 | Synthetic Fur adoption rate in final year (Year 10) | 0.98 |  |  | Input assumption | Target adoption after sustained outreach |  |  |  |
| C5 | Maintainance Cost Escalation | 0.05 |  |  | Input assumption | Expected real wage & logistics growth (5 %) |  |  |  |
| Financial Year | No. Year | Synthetic Adoption rate | Synthetic durability proportion | Mean Effective leopards prevented | Min Effective leopards prevented | Max Effective Leopards prevented | Maintenance cost (nominal) in USD | Discount factor | Discounted maintenance cost in USD |
| 2019-2020 | 1 | 0.65 | 1 | 28.6764705882353 | 16.25 | 182.8125 | 0 | 1 | 0 |
| 2020-2021 | 2 | 0.686666666666667 | 1 | 30.2941176470588 | 17.1666666666667 | 193.125 | 37589.26 | 1 | 37589.26 |
| 2021-2022 | 3 | 0.723333333333333 | 1 | 31.9117647058824 | 18.0833333333333 | 203.4375 | 39468.723 | 1 | 39468.723 |
| 2022-2023 | 4 | 0.76 | 1 | 33.5294117647059 | 19 | 213.75 | 41442.15915 | 1 | 41442.15915 |
| 2023-2024 | 5 | 0.796666666666667 | 1 | 35.1470588235294 | 19.9166666666667 | 224.0625 | 43514.2671075 | 1 | 43514.2671075 |
| 2024-2025 | 6 | 0.833333333333333 | 1 | 36.7647058823529 | 20.8333333333333 | 234.375 | 45689.980462875 | 1 | 45689.980462875 |
| 2025-2026 | 7 | 0.87 | 1 | 38.3823529411765 | 21.75 | 244.6875 | 47974.4794860188 | 0.925925925925926 | 44420.8143389063 |
| 2026-2027 | 8 | 0.906666666666667 | 1 | 40 | 22.6666666666667 | 255 | 50373.2034603197 | 0.857338820301783 | 43186.9028294922 |
| 2027-2028 | 9 | 0.943333333333333 | 1 | 41.6176470588235 | 23.5833333333333 | 265.3125 | 52891.8636333357 | 0.79383224102017 | 41987.2666397841 |
| 2028-2029 | 10 | 0.98 | 1 | 43.2352941176471 | 24.5 | 275.625 | 55536.4568150025 | 0.735029852796453 | 40820.9536775678 |
|  |  |  | Totals | 359.558823529412 | 203.75 | 2292.1875 | 414480.393115052 |  | 378120.327206125 |
| D1 | Effective leopards prevented from ceremonial skin use | 359.558823529412 | 203.75 | 2292.1875 |  |  |  |  |  |
| D2 | Nominal cost per leopard use prevented in USD | 2024.81025488031 | 317.617294883185 | 3573.19456743584 |  |  |  |  |  |
| D3 | Discounted cost per leopard use prevented in USD | 1923.68614519495 | 301.754689442345 | 3394.74025622638 |  |  |  |  |  |
