## Supplementary Material 4 for "Synthetic Substitutes as a Conservation Tool: Evaluating Synthetic Leopard Fur for Demand Reduction and Species Recovery"

Supplementary Material S4: Integrated Mechanism of Change Evidence Details

| **Intervention** | **Expected Outputs** | **No.** | **Evidence** | **Conclusion** |
| --- | --- | --- | --- | --- |
| Saving Spots | Royal and prospective paddlers are aware of synthetic furs | 1.1 | 81.9 % of the 279 paddlers survyed across 2018-2024 were aware of synthetic furs | Saving Spots program led to the decline in acquisition of authentic leopard furs by Lozi paddlers |
|  | Royal and prospective paddlers have access to synthetic furs | 1.2 | Synthetic furs in sufficient quantity were provided to the BRE and distributed to the paddlers before the Kuomboka |  |
|  | Royal and prospective paddlers have a positive perception of synthetic furs | 1.3 | Pooled across years, 78.5 % of respondents (215 of 276) expressed a positive opinion of synthetic furs |  |
|  | Decrease in desire for acquiring authentic leopard skin among paddlers | 1.4 | Only 21 % of paddlers wished to obtain a real skin. The desire to acquire authentic skills declined by |  |
|  | Increase in adoption of synthetic fur | 1.5 | The use of authentic skins during the Kuomboka ceremony plummeted from 2020 to 2024. 39% of royal paddlers wore authentic skins in 2022, compared with just 10% in 2024, representing a 74.4% decline over two years |  |
|  | Decrease in acquisition and ownership of authentic leopard fur | 1.6 | Actual ownership, however, declined sharply by 77.6% reduction from 2020 to 2024.. In 2020, 67 % of paddlers (57 of 86) possessed at least one leopard skin; by 2024 this had fallen to 15 % (coefficient = –0.59, p < 0.01). |  |
| Counter-Trafficking Operations in Western and Southern Zambia | Increase in wildlife-related cases detected and people arrested. | 2.1(a) | The number of reported wildlife-related cases increased from 80 in 2018 to 213 in 2023 — an overall increase of 166%. This corresponds to an average annual growth rate of 20.8% over the five-year period. In 2024, the number of cases dropped to 139 - a decrease of 34.7% compared to 2023. | Counter-trafficking operations disruped illegal wildlife trade in Southern and Western Zambia |
|  |  | 2.1(b) | The number of people arrested in wildlife-related cases from 2018 to 2023 increased from 117 to 232, representing a nearly 98% overall rise, with an average annual growth rate of around 14.5%. In 2024, 137 people were arrested representing a drop of about 41% compared to 2023. |  |
|  | Offenders are prosecuted in court and convicted. | 2.2 | Between 2018 and 2024, the cumulative number of individuals arrested for wildlife-related offenses increased from 117 to 1,146, while cumulative convictions secured rose from 80 to 867. The cumulative conviction rate increased from 68% in 2018 to over 75% by 2024 reflecting a high conviction rate and steady improvement. |  |
|  | Offenders are incapacitated | 2.3 | No. offenders detected from 2018 to 2024 were re-arrested in wildlife-related cases during the same period. This may indicate low levels of re-offending among arrested poachers. |  |
|  | Increased perception of risk of arrests among traffickers. | 2.4 | *No information available |  |
|  | Potential offenders deterred | 2.5 | *No information available |  |
|  | Decline in trafficking of leopards | 2.6 | The number of wildlife-related cases involving leopards declined significantly from 11.25% of all cases in 2018 to 2.88% in 2024 (Result 2.1) |  |
| Counter-Poaching patrols inside GKE | Improved patrol vigilance in GKE | 3.1(a) | Patrol Effort increased by XX % from YYYY kms in 2018 to ZZZZ kms in 2024. | Anti-poaching patrols improved vigilance resulting in decline in poaching intrusions in the park. |
|  |  | 3.1(b) | Patrol Coverage increased from XX % in 2018 to YY % in 2024 |  |
|  | Increase in detection of poachers | 3.2 | No information patrol data available prior to SMART introduction. But anectodal sources suggest SMART improved detection. |  |
|  | Increase in detection of snares and traps | 3.3 | No information patrol data available prior to SMART introduction. But anectodal sources suggest SMART improved detection. |  |
|  | Increase in risk perception of patrols among poachers | 3.4 | *No information available. |  |
|  | Increased effort for poachers to poach leopard prey or leopards in Kafue | 3.5 | *No information available. |  |
|  | Poachers dettered from poaching in Kafue | 3.6 | *No information available. |  |
|  | Decrease in poaching activity in the park. | 3.7(a) | Encounter rate of snares and traps (snares and traps/ 100 km patrol) decreased from 2018-2024 |  |
|  |  | 3.7(b) | Encounter rate of poacher apprehensions and human activity decreased in the park from 2018 to 2024. |  |
